# Neural interference between real and imagined visual stimuli

**DOI:** 10.1101/2024.01.05.574285

**Authors:** Alexander A Sulfaro, Amanda K Robinson, Thomas A Carlson

## Abstract

Evidence suggests that mental imagery and veridical perception recruit similar components of the human visual system. If so, neural representations of imagined and real stimuli should interact with one another, combining constructively or competing antagonistically. To determine if and how real and imagined visual stimuli interact in the brain, we asked participants to mentally visualise white bars at specific orientations after a rhythmic countdown while their brain activity was recorded using electroencephalography. Stimuli were imagined in isolation, or while another stimulus at a highly or poorly congruent orientation appeared on-screen. Multivariate pattern analysis was used to assess whether overlap between imagined and real stimulus features enhanced or diminished stimulus-specific sensory information in the brain. Findings showed that imagined and real orientation could be decoded from brain activity, with real orientation decoding mildly amplified by highly congruent, but not poorly congruent, imagined orientations. Although interactions between real and imagined stimuli were observed, no evidence was detected to suggest that imagined and real stimuli use the same neural activity patterns to encode sensory information. Instead, congruent imagery seemed only to amplify activity which had already been induced by real percepts, targeting late- but not early-stage perceptual representations. Ultimately, this study suggests that imagined and real stimuli interact in a mildly constructive manner, with imagination mostly acting in a modulatory capacity.

## Introduction

Mental images are created during waking life, seemingly while our brains are simultaneously processing sensory inputs from the external environment. Yet, real and mental image perception both seem to demand similar neural resources (Albers et al., 2013; Bosch et al., 2014; Cichy et al., 2012; Dijkstra et al., 2017, 2018, 2019, 2020; Lawrence et al., 2018; Lee et al., 2012; Naselaris et al., 2015; Xie et al., 2020). Generating a mental image should therefore be expected to interfere, compete, or in some way interact with veridical processing of the external world. Such interference may even account for why mental images appear as quasi-sensory percepts, unlike real percepts, hallucinations, and dreams (Sulfaro et al., 2023). However, the specific manner in which imagined content interacts with veridical content is yet to be determined.

If imagined and veridical perception rely on shared neural representations, it might be expected that they add together constructively. Accordingly, mental images tend to prime veridical images of similar content in binocular rivalry tasks (Keogh & Pearson, 2011, 2014, 2017; Pearson et al., 2008; Sherwood & Pearson, 2010). Mental imagery also lowers the detection threshold for low-contrast veridical stimuli (Dijkstra et al., 2021, 2022). This could suggest that imagined and real percepts share the same neural code, amplifying signals from one another, with imagined stimuli simply activating perceptual representations to a weaker extent than real ones (Koenig-Robert & Pearson, 2021; Pearson et al., 2015). Conversely, if imagined and veridical perception rely on the same representational resources in the brain, yet use distinct neural codes, they might be expected to destructively compete against or detract from one another. Under this view, each type of percept should antagonise the persistence of the other. Studies have shown that increasing the luminance of environmental stimuli during the generation or maintenance of mental imagery can reduce the priming effects of mental imagery in binocular rivalry tasks (Keogh & Pearson, 2011; Pearson et al., 2008; Sherwood & Pearson, 2010). The recall of modality-specific information is also impeded by the appearance of real distractors in the same sensory modality (Vredeveldt et al., 2011; Wais et al., 2010), and one study showed greater impairment for distractors with more similar image statistics to the imagined stimulus (Borst et al., 2012). Overall, evidence regarding the nature of real-imagined interactions seems to be mixed and generally relies on indirect behavioural evidence such that the specific nature of real-imagined stimulus interactions remains an open question. Consequently, here we sought to determine whether real-imagined interactions generally interact in a constructive way, or in a destructive way.

Assessing the nature of interference between real and imagined content in the brain is complicated by the lack of neural measures indexing the quality of a mental image. However, previous works have proposed that mental image quality might be understood as the degree to which a thought gains sensory properties in addition to abstract properties (Sulfaro et al., 2023). Although the relationship between neural processes and imagined experience is far from understood, a measure that quantifies stimulus-specific information in the brain during imagery can at least indicate how much sensory information is available for imagination to use. Neural decoding and multivariate pattern analysis (MVPA) offer such an approach (Dijkstra et al., 2018; Grootswagers et al., 2017; Linde-Domingo et al., 2019; Robinson et al., 2021). Neural decoding involves training a machine-learning classifier to extract regularities in brain activation patterns associated with specific events, such as seeing or imagining a particular image. When applied to novel brain data, a trained classifier can make predictions about which specific event induced a given pattern of brain activity. When classifying basic sensory features of real or imagined stimuli, the accuracy of classification should depend on how much sensory information is present in the brain to discriminate between different stimuli. Changes in the accuracy of imagined or real feature decoding might therefore be one way of judging whether the real and imagined features constructively or destructively interact with one another.

As visual mental image content is carried via feedback signals (Dentico et al., 2014; Dijkstra et al., 2020; Linde-Domingo et al., 2019), feedforward interference could disrupt or erase mental image representations on a short timescale given that the visual system oscillates between feedforward and feedback processing sweeps at an alpha-band frequency (Dijkstra et al., 2020). Temporally resolved decoding is therefore most optimal for investigating interference effects between real and imagined stimuli, using electroencephalography (EEG) or magnetoencephalography (MEG). Yet, previous studies attempting time-resolved decoding of mental images have generally relied upon complex stimuli, such as faces, objects, and scenes (Dijkstra et al., 2018; Linde-Domingo et al., 2019; Shatek et al., 2019). Classifiers trained to decode complex imagined stimuli such as these could capitalise on neural patterns relating to the semantic content, rather than the sensory content, of these imagined stimuli. For instance, one study explored the low-level features of mental images by decoding between imagined photographs and imagined drawings (Linde-Domingo et al., 2019), yet this distinction could still be reduced to a high-level categorical distinction. While it is unclear if the visual features of a stimulus can ever be truly divorced from its semantic features, studying the most basic visual features of imagined content possible should improve the likelihood that imagined stimulus information decoded from brain activity is sensory in nature.

Previous studies have also had mixed success with regard to decoding the contents of mental images using EEG during periods when imagery was expected (Linde-Domingo et al., 2019; Shatek et al., 2019). In part, this may be because mental image perception tends to generate less neural activity than real image perception (Bainbridge et al., 2021; Dijkstra et al., 2017, 2018; Harrison & Tong, 2009; Lee et al., 2012; Reddy et al., 2010; Robinson et al., 2021), and because mental images are less temporally precise given that they are unobservable, self-generated events. To improve the temporal precision of imagery, one study had participants track discrete movements of an imagined shape in sequence with a rhythmic timing cue, allowing the successful decoding of imagined position using EEG (Robinson et al., 2021). Rhythmic countdowns have also been used to successfully study imagined sounds using EEG (Jack et al., 2019; Whitford et al., 2017). Rhythmically cuing imagery should therefore improve the temporal consistency of visual imagery across trials, allowing a machine learning classifier to make a better estimate of the neural activity patterns associated with imagined content at specific times.

Here, we describe a study using neural decoding to investigate the temporal dynamics of imagined stimulus representations with and without interference from real visual stimuli. Neural activity was recorded with 128-channel EEG while participants were imagining simple bar stimuli at different orientations. EEG was used as a non-invasive, accessible, and temporally precise method of recording brain activity. In order to ascertain the nature of real-imagined stimulus interactions in the brain, our study compared how easily sensory information could be decoded when imagined and real stimuli were highly congruent in orientation compared to poorly congruent. To reduce the variability in the onset of mental image generation, participants were cued to produce a mental image following a rhythmic countdown. Orientation was decoded from imagined stimuli as it is a basic visual feature which may be difficult to recall without a visual mental imagery strategy. Hence, imagined orientation decoding in this study is presumed to index sensory, rather than abstract, information in neural representations. If imagined and real stimuli interact constructively, decoding which orientation was viewed or imagined should be most accurate when imagined and real stimuli are highly congruent. However, if imagined and real stimuli compete against one another, then orientation decoding should be most accurate when real and imagined stimuli are dissimilar.

## Methods

The experiment consisted of two sections, with EEG recorded throughout both. In the first part (mental imagery trials), participants were sequentially presented with two bars of different orientation, and then cued to construct a mental image of one of the bars at a specific time in the presence or absence of a bar stimulus presented on-screen. Data collected during this task was used to analyse the neural representations of imagined content and their interactions with veridical perceptual content. In the second section (passive viewing trials), participants passively viewed bars of different orientations. EEG recordings during this phase were used to estimate the patterns of neural activity associated with the veridical perception of orientation.

### Participants

Undergraduate psychology students at The University of Sydney were recruited to participate in the study in exchange for course credit. The study was approved by the university ethics committee. Students were excluded from participating if they had a history of head injury, neurological, or psychiatric illness, if their body contained non-removable metal, or if they did not have normal or corrected-to-normal vision. In total, 51 participants gave informed consent to participate. 3 participants were excluded from analysis after they reported imagining stimuli which deviated by ≥22.5° degrees on average from the expected orientation during memory checks. Of the remaining 48 participants, 13 were male and 35 female, with a median age of 19 (range 17-32). 44 were right-handed and 4 were left-handed.

### Experimental materials

Stimuli were presented on an Asus VG236H monitor with a 60Hz refresh rate and 95.79 PPI (1920×1080 resolution). Responses were recorded via button press on a Dell KB212-B keyboard. Stimulus creation and the experimental task was programmed using custom Python code and the PsychoPy library (Peirce et al., 2019). Visual image presentations were confirmed to deviate from expected timings by no more than 6ms on average using Black Box Toolkit photodiode timing tests.

### Stimuli

Each stimulus consisted of a single white bar, 7° of visual angle long, viewed against a black background. Each bar was created by filtering a 0.005Hz spatial frequency cosine wave through a raised cosine masking function to produce a single bar blurred at the edges. A simple bar of low spatial frequency was chosen to increase the likelihood of interference between real and mental images given that imagination invokes lower spatial frequency representations relative to during veridical perception (Breedlove et al., 2020). All stimuli were viewed in a pitch-black room, centred on a computer screen, with screen brightness reduced to the lowest level to reduce discomfort. The bars participants were asked to visualise were orientated at either 292.5°, 337.5°, 22.5°, or 67.5° from vertical.

Interfering bars which appeared on-screen during imagery deviated from the imagined orientation by ±22.5° (highly congruent interference) or ±67.5° (poorly congruent interference) such that interference bars were oriented at either 315°, 0°, 45°, or 90° from vertical.

A timing marker was used to indicate when imagery should occur. This marker consisted of four red triangles pointing inwards, each centred at the top, bottom, left and right of the screen, 5° of visual angle from the screen centre. This marker was also on-screen during the presentation of all real stimuli. A white version of the marker also flashed on-screen to provide aid as a rhythmic countdown to the imagery period.

Post-stimulus masks during imagery trials consisted of 100 2Hz, Gaussian-masked, black-and-white gratings of 4° of visual angle, randomly oriented within a 7° field. Grating positions were randomly sampled from a normal distribution with a mean in the centre of the screen. Masks were generated dynamically on each trial.

### Procedure

#### Mental imagery task

For the mental imagery task (Figure 1), on each trial, participants were sequentially presented with two stimuli and then asked to visualise one of them. Participants were required to imagine only one of the two stimuli such that the other could be used as an unimagined control. Trials began with a grey fixation cross presented in the centre of the screen for 1000ms, followed by an oriented bar stimulus for 800ms, then an 800ms random-grating mask. This sequence was repeated for the second stimulus orientation to be encoded. After the second sequence, a fixation cross was presented for 800ms before the number 1 or 2 appeared for 800ms, instructing the participant that they would be imagining the first or second orientated stimulus. Before constructing a mental image, participants were presented with a rhythmic flashing countdown to help them precisely time the onset of their mental imagery. This countdown consisted of three repeated instances of a 500ms fixation cross, followed by a 100ms presentation of a timing marker: four white triangles in the periphery, pointing inwards. After the third flash of the timing marker and another 500ms fixation period, the timing marker appeared again for 500ms in red to demarcate the period in which mental imagery should occur.

**Figure 1.**
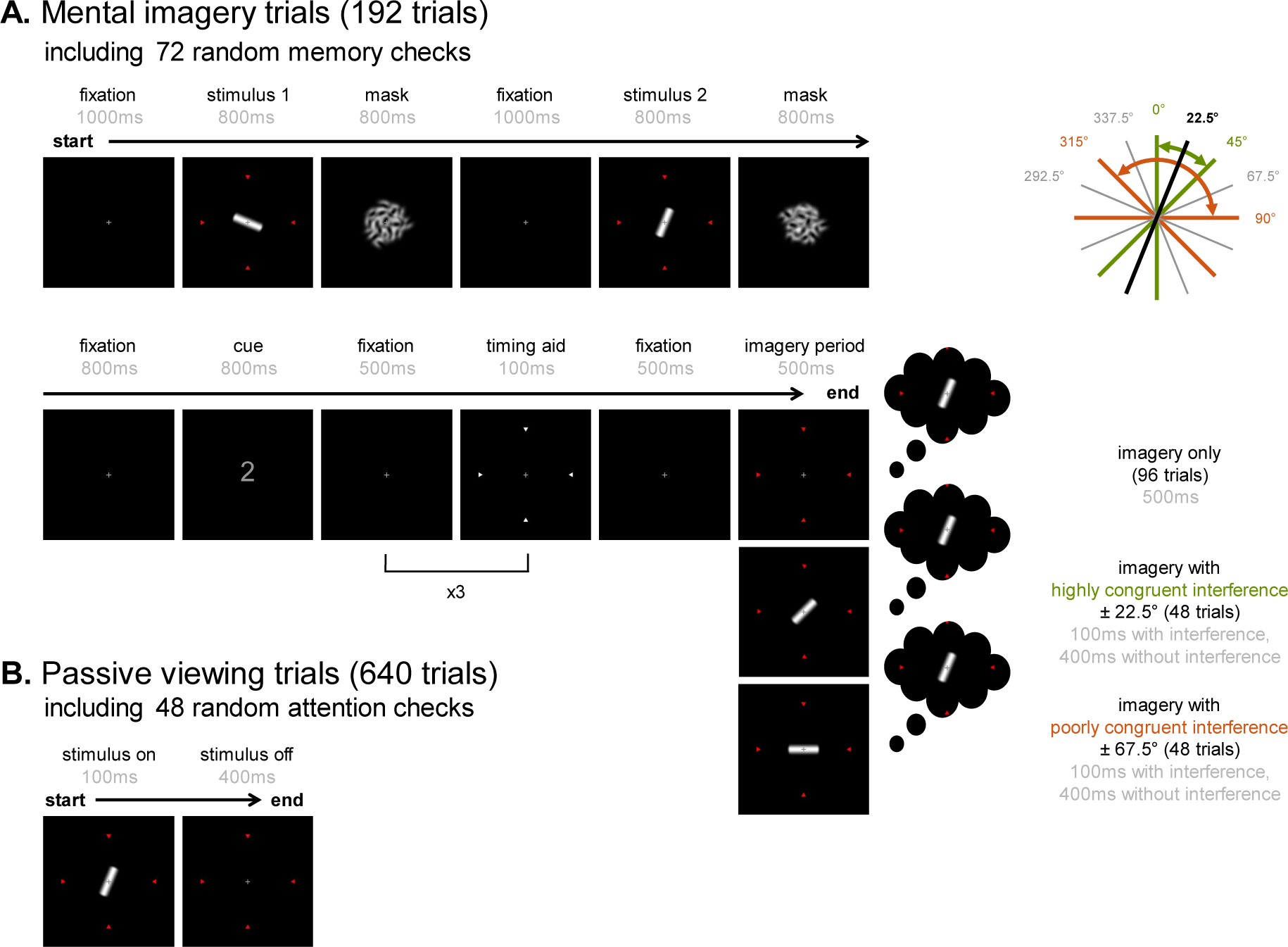
Experimental procedure for imagery and passive viewing trials. **A.** Mental imagery trials involved encoding two sequentially presented stimuli of different orientations (shown in black/grey, top right). A numerical cue indicated which stimulus was to be visualised during the imagery period. Four white markers flashed rhythmically on-screen before the imagery period to help participants time their mental images precisely. Another bar stimulus appeared on-screen during the first 100ms of the imagery period for half of imagery trials. This interfering stimulus was either closely oriented to the imagined stimulus (highly congruent), or distally oriented (poorly congruent). Possible orientations for the interfering stimulus are shown in green and orange (top right), coloured according to their congruency relative to the cued orientation in this example. Participants were occasionally required to reconstruct the orientation they imagined at the end of trials to check their recall accuracy. **B.** Passive viewing trials involved viewing a continuous stream of different orientations separated by fixation screens. Participants pressed a button whenever the central fixation cross turned red, which happened on random trials. All eight orientations used in the experiment were shown during passive viewing.

Half of the trials involved an imagery period with no other stimulus present on-screen (no interference), and the other half involved an imagery period with a concurrent stimulus presented on-screen (interference). Three scenarios were possible: imagery without interference, imagery with highly congruent interference, or imagery with poorly congruent interference. During highly congruent interference, a white bar with an orientation ±22.5° from the imagined orientation would appear. During poorly congruent interference, a white bar with an orientation ±67.5° from the imagined orientation would appear. When interference was present, it lasted for the first 100ms of the 500ms imagery period.

There were 6 blocks of 32 imagery trials, with 192 imagery trials in total. Each block contained 8 instances of imagery per orientation: 4 instances without interference, 2 with highly congruent interference, and 2 with poorly congruent interference. The order of imagery trials was randomised within a block, and the sequential order in which bars were encoded, and cued for recall, was also randomised and counterbalanced across the experiment. Each trial automatically followed the previous trial, with a blinking fixation cross to indicate the start of a new trial. Breaks were provided at the midpoint and end of each block.

To verify that participants were trying to construct a mental image with the expected orientation, each block also had 12 trials with memory checks where participants were required to reconstruct the orientation of the cued stimulus by using the arrow keys to rotate a bar on-screen. Only the endpoints of the bar were visible to facilitate easier reimagining of the stimulus if required. Participants were given 20 seconds to respond, with a countdown timer in the lower-right of the screen. To incentivise performance and gamify the task, participants were given immediate qualitative feedback on their check-trial response accuracy: “Perfection!” for ≤1° of deviance, “Exceptional!” for up to 10° of deviance, “Good!” for up to 20°, “Okay” for up to 30°, and “Not quite” for >30°. Average deviances were given as feedback at the end of each block. Each block contained 3 memory checks for each of the 4 possible imagined orientations: 1 for trials without interference, 1 for trials with highly congruent interference, and 1 for trials with poorly congruent interference. There were 3 memory checks for each possible combination of imagined and interfering angle over the experiment.

#### Passive viewing (pattern estimator)

After the completion of all mental imagery trials, participants engaged in a short task where they passively viewed a stream of oriented white bars. This task was included to allow us to estimate the neural signatures associated with externally generated stimulus perception. Passive viewing trials involved alternating presentations of the oriented bar stimuli used in imagery trials shown on-screen for 100ms, followed by a fixation cross for 400ms, replicating the timing of interference during imagery trials. Each subsequent passive viewing trial continued immediately after the previous. To maintain attention, participants were required to press a button as quickly as possible whenever they noticed the fixation cross turning from grey to red, which occurred pseudorandomly, and for the entirety of a 500ms trial.

There were 20 blocks of 32 passive viewing trials, with 640 on-screen stimulus presentations in total. In each block, there were 4 presentations of each of the 4 orientations cued during imagery trials and for each of the 4 interfering orientations. In total, there were 6 response trials with a red fixation cross for each of the 8 unique orientations. Response trials were randomly distributed across the whole session and were not analysed.

### EEG recording and pre-processing

EEG was recorded using actiCAP and actiCHamp Brain Vision systems (Brain Products GmbH). Electrical potentials were sampled continuously at 1000Hz from 128 electrodes. Electrodes were positioned according to the 10-5 system (Oostenveld & Praamstra, 2001) with an online reference electrode at location FCz. After the recording session, EEG data was minimally pre-processed (Grootswagers et al., 2017) using MNE-Python (Gramfort et al., 2013). Electrode measurements were re-referenced to the average of all electrodes. A 50Hz notch filter was then applied to remove line noise, followed by a 100Hz lowpass filter and 0.1Hz highpass filter before downsampling to 250Hz. The continuous recording was segmented into epochs ranging from −500ms to 1000ms around real image onset and/or expected mental image onset. Data was analysed both with all 128 channels and with a 56-channel subset representing the posterior portion of the head: CPz, CP1, CP2, CP3, CP4, CP5, CP6, Pz, P1, P2, P3, P4, P5, P6, P7, P8, POz, PO3, PO4, PO7, PO8, Oz, O1, O2, TP7, TP8, TP9, TP10, TPP9h, TPP7h, CPP5h, CPP3h, CPP1h, CPP2h, CPP4h, CPP6h, TPP8h, TPP10h, P9, PPO9h, PPO5h, PPO1h, PPO2h, PPO6h, PPO10h, P10, PO9, POO9h, POO1, POO2, POO10h, PO10, O9, OI1h, OI2h, and O10. Results for each 128-channel analysis are shown in the main text, with the corresponding 56-channel analysis shown in Appendix 1.

### Neural decoding

Neural decoding was carried out to evaluate the degree to which stimulus-specific information about imagined and real stimuli is embedded in neural activity patterns. If stimulus-specific information can be successfully extracted, decoding can be used to investigate the nature of interference between real and imagined stimuli.

Decoding accuracy over time was assessed both for real and imagined orientations in each condition. For periods of real or mental image perception, a machine-learning classifier was trained for each participant for each timepoint of the epochs of EEG data recorded during a condition. Timepoint-by-timepoint classification allowed the temporal dynamics of imagined and real feature representations to be assessed. Classifiers were trained using linear discriminant analysis (LDA) from the scikit-learn Python package (Pedregosa et al., 2011) using each EEG channel as a feature, and voltage recordings during each trial as samples. For orientation decoding, classifiers were trained to decode the orientation of a real or imagined stimulus in a pairwise manner (e.g. 22.5° vs. 67.5°), with a separate classifier trained and tested for every unique pairwise combination of angles for a given condition. Chance-level decoding accuracy was therefore 50% for all analyses. Decoding accuracy was determined by calculating the proportion of test trials correctly classified by a trained classifier. Overall pairwise decoding accuracy was determined by taking the average of the individual pairwise decoding accuracies, then averaging across participants. Where the classifier was trained and tested using the same dataset, a leave-one-block-out cross-validation structure was used (6-fold for imagery trials, 20-fold for estimator trials).

### Statistical inference

Bayes factors (BFs) were calculated to determine whether decoding accuracy was meaningfully above chance for each timepoint, and if decoding differed between conditions (Rouder et al., 2009; Teichmann et al., 2022). Bayes factors reflect the probability of the observed data under the alternative hypothesis (above-chance decoding, or a difference in decoding) relative to the null hypothesis (chance-level decoding, or no difference in decoding). Bayesian t-tests were conducted using Cauchy priors with a scale factor of 0.707 and an effect size of 0.5, following the method used by Teichmann et al. (2022). One-tailed priors were used to assess greater-than-chance decoding accuracy as below-chance decoding accuracy is not meaningfully interpretable here. Two-tailed priors were used to assess differences in decoding accuracy between conditions. Bayes factors of 3 or greater indicate some evidence in favour of the alternative hypothesis, with strong evidence indicated by Bayes factors of 10 or greater. Bayes factors of 1/3 or below indicate evidence in favour of the null, with values between 1/3 and 3 reflecting insufficient evidence for either hypothesis.

## Results

### Behavioural analysis (recall accuracy)

Of the 192 mental imagery trials, 72 required participants to reconstruct the orientation of the stimulus they had been cued to imagine. For the 48 included subjects, collapsing across conditions, orientation reconstructions deviated from the cued orientation by a mean (M) magnitude of 11.5° with a standard deviation (SD) of 4.4°. Paired t-tests revealed no significant difference in absolute reconstruction error between imagery trials with on-screen interference (M = 11.7°, SD = 4.7°) and those without interference (M = 11.2°, SD = 4.6°; t(47) = 1.2, p = .237). However, errors were significantly larger in magnitude in the presence of poorly congruent interference (M = 12.2°, SD = 5.3°) compared to highly congruent interference (M = 11.2°, SD = 4.4°; t(47) = −2.2, p = .033).

### Decoding real orientation

Neural responses to real stimuli tend to be more prominent than to imagined stimuli (Bainbridge et al., 2021; Dijkstra et al., 2017, 2018; Harrison & Tong, 2009; Lee et al., 2012; Reddy et al., 2010; Robinson et al., 2021). To confirm that orientation was possible to decode from brain activity for real stimuli, average pairwise decoding accuracy was determined over time for oriented stimuli seen during passive viewing, and for oriented stimuli which appeared on-screen as interference during imagery trials. Average pairwise classification accuracy over time for real orientations is shown in Figure 2A. Above-chance decoding suggested that orientation representations associated with real stimuli could be extracted from both passive viewing and imagery trials.

**Figure 2.**
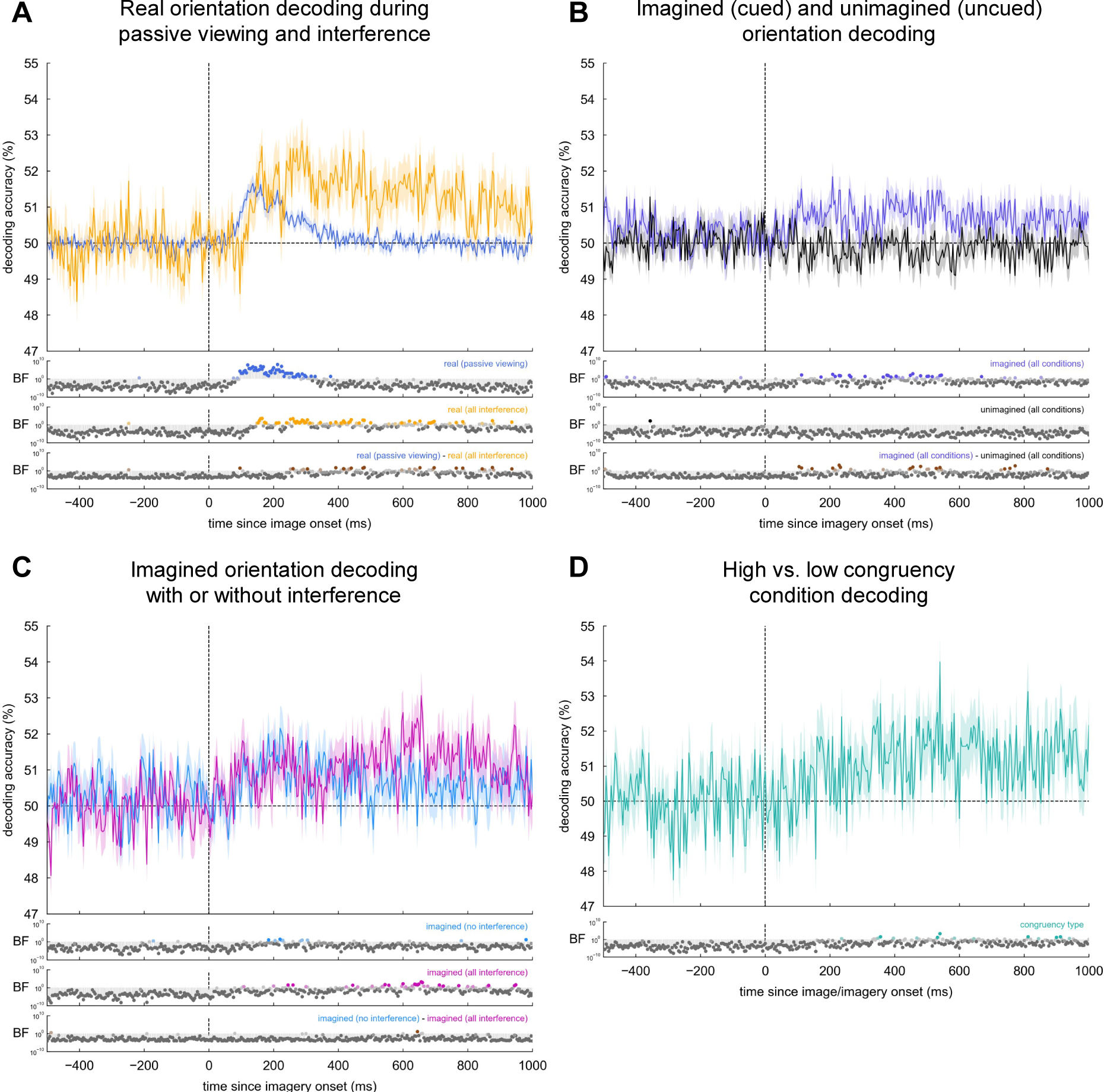
Decoding of real orientation, imagined orientation, and congruency type (128 channels) Decoding accuracy over time from 128-channel EEG recordings. Chance decoding accuracy is 50%. Bayes factors (BF) are shown for the alternative hypothesis of above-chance decoding, or for a difference in decoding accuracy between two conditions. BF ≥ 10 (saturated colour), 3 ≤ BF < 10 (pale colour), 1/3 ≤ BF < 3 (light grey), 1/10 ≤ BF < 1/3 (grey), BF < 1/10 (dark grey). **A.** Decoding the orientation of real stimuli presented on-screen, either during passive viewing trials, or during interference trials with concurrent mental imagery. **B.** Decoding the orientation of the stimulus cued to be imagined in the imagery period (imagined), and of the stimulus which was encoded but not recalled (unimagined). **C.** Decoding imagined orientation when no competing stimulus was on-screen (no interference), or when a competing stimulus of any orientation was on-screen at the same time (all interference). **D.** Decoding whether an imagery trial was presented at the same time as a highly or poorly congruent stimulus on-screen. See Appendix 1.2 for 56-channel analyses using only posterior electrodes.

### Decoding imagined orientation

Neural decoding was carried out to determine if there was sufficient information in the brain to discriminate between imagined orientations. Note that classifying imagined orientation relies on fewer orientation samples than when classifying real orientation in this experiment. Signal-to-noise ratio is therefore reduced when classifying imagined orientations.

Despite encoding two orientations per imagery trial, participants were only cued to imagine one orientation during the imagery period. Significant above-chance decoding accuracy was obtained for the cued (imagined) orientation, but not the uncued (unimagined), indicating that classification during the imagery period was not merely due to lingering neural activity from encoding the real cued or uncued stimulus earlier. Imagined and unimagined orientation decoding is shown in Figure 2B.

Half of imagery trials involved interference, involving a real bar appearing on-screen during the first 100ms of the imagery period. Average pairwise decoding of imagined orientation is shown for imagery during interference and for imagery without any interference in Figure 2C. Information about the imagined orientation was present ∼200ms after imagery began in both cases, although interference tended to extend the period of time in which imagined orientation could be decoded. However, there were almost no time points in which evidence favoured a difference in decoding accuracy between the interference and no-interference conditions.

### Decoding congruency condition

Before testing the specific impact of high- or low-congruency interference on orientation decoding, classifiers were trained and tested at each time point to decode whether an interference trial occurred during high- or low-congruency imagery. Each congruency condition involved the same real and imagined stimuli, but experienced in different combinations. Above-chance decoding of congruency condition therefore indicates if the *relationship* between real and imagined stimuli was encoded in the brain, regardless of if imagined and real stimuli interact in any substantial, stimulus-specific manner. Classification between high- and low-congruency conditions was above chance following stimulus onset when analysing the 128-channel dataset, although evidence in favour of an above-chance effect was present only for a handful of timepoints (Figure 2D).

### Orientation decoding modulated by real-imagined congruency

When interference occurred during imagery trials, real stimuli were presented on-screen at an orientation with either high or low congruency to the imagined orientation. Real-imagined congruency did not have any notable impact on the ability of the classifier to decode the orientation of imagined bars (Figure 3B). However, congruency had a mild impact on decoding the orientation of real bars (Figure 3A), with highly congruent interference leading to improved decoding accuracy relative to poorly congruent interference, largely consistent with a constructive interaction between real and imagined stimuli. Yet, there was no prolonged evidence for a difference in decoding accuracy between the congruency conditions.

**Figure 3.**
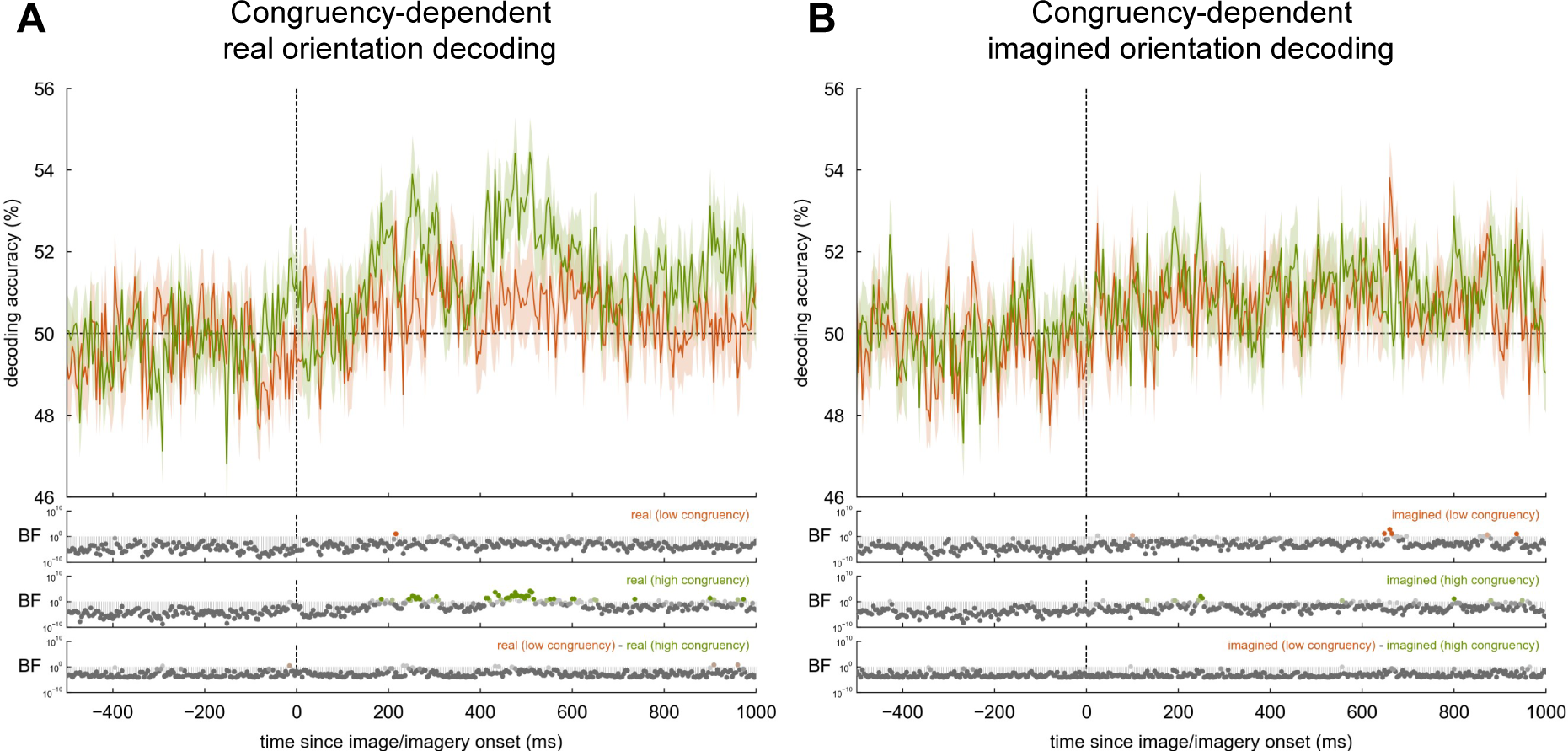
Orientation decoding during high and low congruency interference conditions (128 channels) Decoding accuracy over time from 128-channel EEG recordings. Chance decoding accuracy is 50%. Bayes factors (BF) are shown for the alternative hypothesis of above-chance decoding, or for a difference in decoding accuracy between two conditions. BF ≥ 10 (saturated colour), 3 ≤ BF < 10 (pale colour), 1/3 ≤ BF < 3 (light grey), 1/10 ≤ BF < 1/3 (grey), BF < 1/10 (dark grey). **A.** Decoding real orientation during poorly congruent mental imagery or highly congruent mental imagery. **B.** Decoding imagined orientation in the presence of a poorly congruent or highly congruent real stimulus. See Appendix 1.3 for 56-channel analyses using only posterior electrodes.

To determine if there was a general effect of congruency on real orientation decoding rather than an effect at any specific timepoint, the area under the curve between the high- and low-congruency real orientation decoding traces was compared for all timepoints after time zero. A Wilcoxon signed-rank test showed that the high-congruency interference condition involved a significantly greater area under the curve compared to the low-congruency condition (Z = −2.8, p = .005), equivalent to an average increase in decoding accuracy of 1.2% per timepoint, but only for the posterior 56-channel subset of data (Appendix 1.3). There was no significant difference in area between the congruency conditions for real orientation decoding using all 128 channels, or for imagined orientation decoding using either 128 or 56 channels (all p > 0.05).

### Time-generalised decoding

To investigate whether congruency between real and imagined stimuli affected the persistence or reactivation of neural representations, time-generalised decoding was also carried out. Classifiers trained at one timepoint can also accurately decode stimulus information from activity patterns at other timepoints if neural representations persist for extended periods or are reactivated at different times. Here, a classifier was trained on activity patterns from each timepoint, then every classifier was tested at every timepoint, resulting in a matrix of decoding accuracies for every combination of training and testing time points. Time-generalised decoding of imagined orientation is shown in Figure 4. Diffuse generalisation was observed, trending towards earlier timepoints where interference was absent and later timepoints where interference was present.

**Figure 4.**
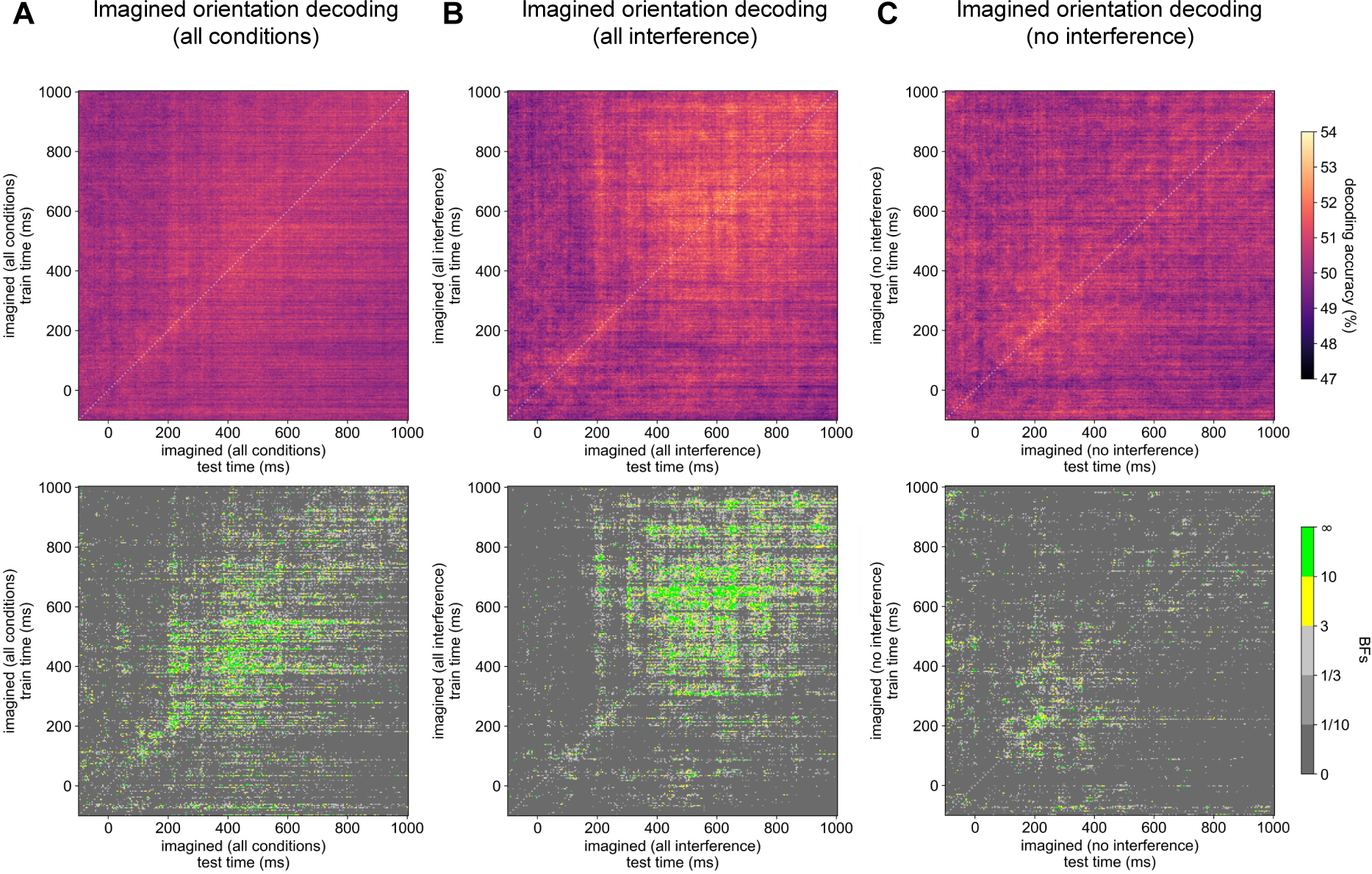
Time-generalised decoding of imagined orientation (128 channels) Time-generalised decoding accuracy, training and testing a classifier at every timepoint, using data from 128-channel EEG recordings. Chance decoding accuracy is 50%. Bayes factors (BFs) are shown for the alternative hypothesis of above-chance decoding. Times are relative to imagery onset. A dotted white line indicates the diagonal. **A.** Time-generalised decoding of imagined orientation for all imagery trials. **B.** Time-generalised decoding of imagined orientation when a competing stimulus of any orientation was on-screen at the same time. **C.** Time-generalised decoding of imagined orientation when no competing stimulus was on-screen. See Appendix 1.4 for 56-channel analyses using only posterior electrodes.

Time-generalised decoding results are shown for the classification of real orientation during high- and low-congruency imagery in Figure 5. When real and imagined orientations were highly congruent, neural activity patterns encoding stimulus information tended to be somewhat persistent, with classifiers trained on earlier timepoints generalising to later timepoints and vice versa. However, little persistent or recurring activity was observed when real and imagined orientations were poorly congruent. Time-generalised classification results are also shown for imagined orientation decoding during high- and low-congruency interference in Figure 6, with classification generalising relatively diffusely in both conditions.

**Figure 5.**
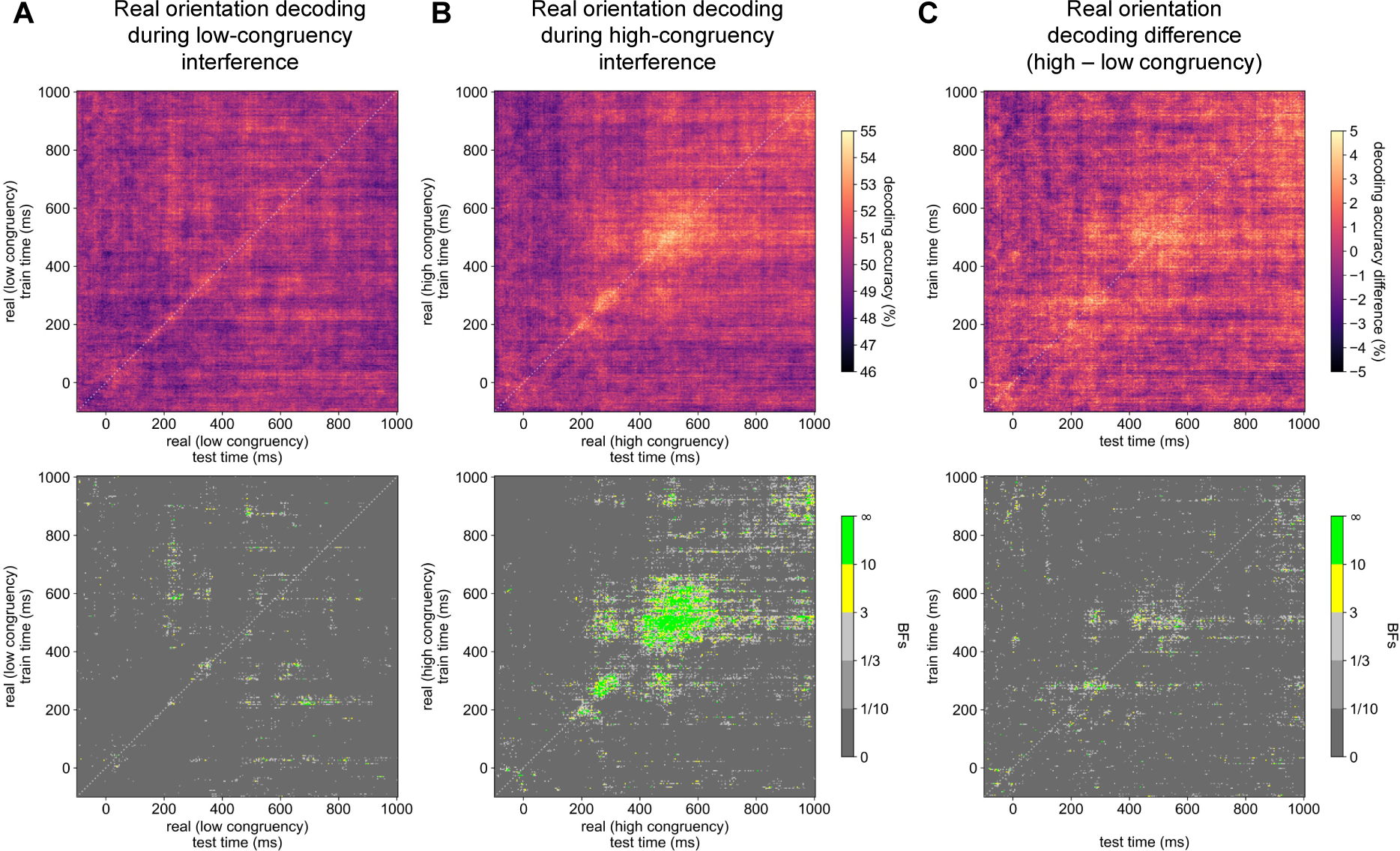
Time-generalised decoding of real orientation dependent on congruency condition (128 channels) Time-generalised decoding accuracy, training and testing a classifier at every timepoint, using data from 128-channel EEG recordings. Chance decoding accuracy is 50%. Bayes factors (BFs) are shown for the alternative hypothesis of above-chance decoding, or for a difference in decoding accuracy between two conditions. Times are relative to image onset. A dotted white line indicates the diagonal. **A.** Time-generalised decoding of real orientation during poorly congruent mental imagery. **B.** Time-generalised decoding of real orientation during highly congruent mental imagery. **C.** The difference in real orientation decoding accuracy between the high- and low-congruency interference conditions. See Appendix 1.5 for 56-channel analyses using only posterior electrodes.

**Figure 6.**
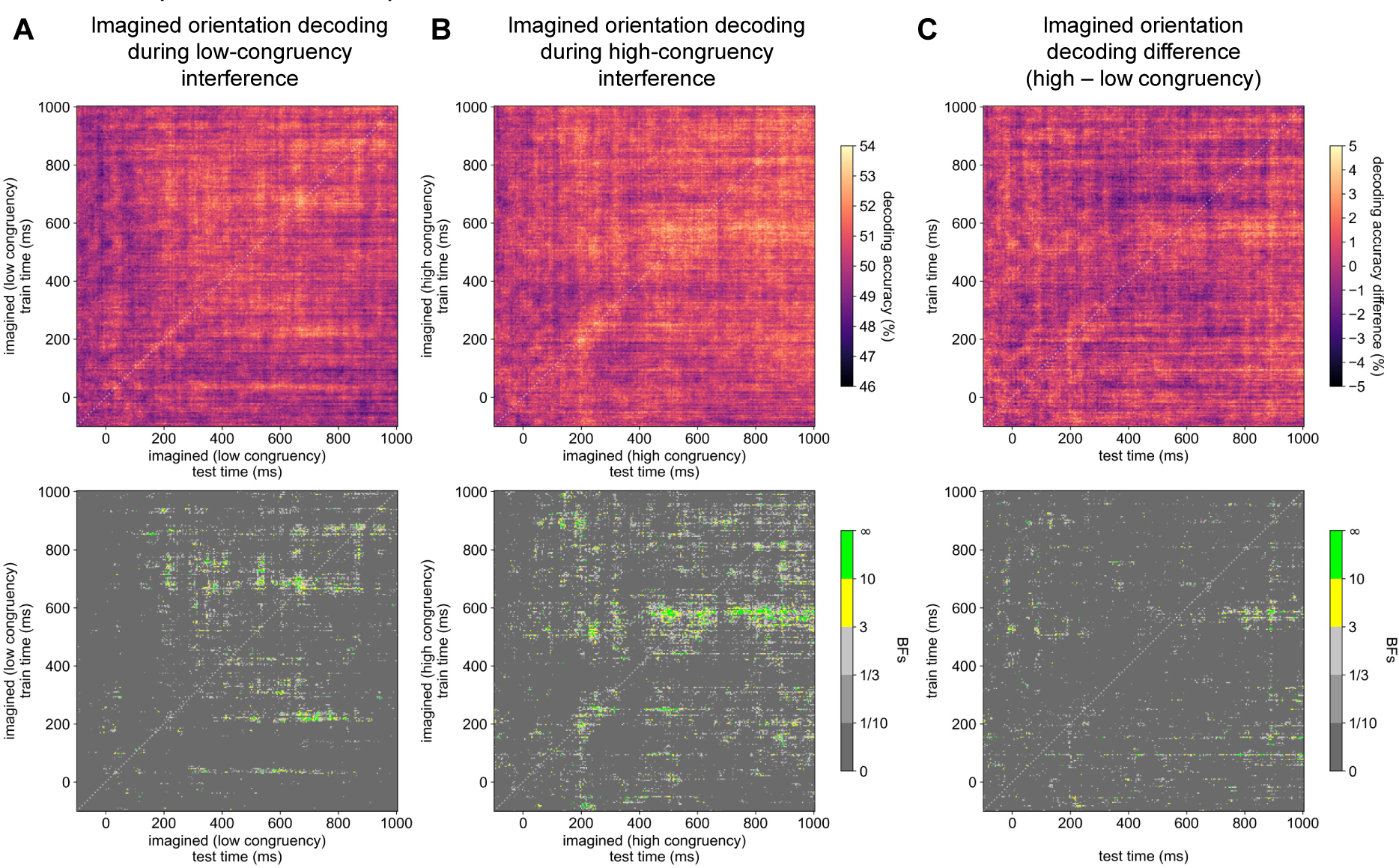
Time-generalised decoding of imagined orientation dependent on congruency condition (128 channels) Time-generalised decoding accuracy, training and testing a classifier at every timepoint, using data from 128-channel EEG recordings. Chance decoding accuracy is 50%. Bayes factors (BFs) are shown for the alternative hypothesis of above-chance decoding, or for a difference in decoding accuracy between two conditions. Times are relative to imagery onset. A dotted white line indicates the diagonal. **A.** Time-generalised decoding of imagined orientation in the presence of a poorly congruent real stimulus. **B.** Time-generalised decoding of imagined orientation in the presence of a highly congruent real stimulus. **C.** The difference in imagined orientation decoding accuracy between the high- and low-congruency interference conditions. See Appendix 1.6 for 56-channel analyses using only posterior electrodes.

### Shared representations between real and imagined stimuli

To investigate the possibility of overlap between low-level real and imagined feature representations, pairwise cross-decoding of orientation was carried out. Cross-decoding involves training a classifier to decode a particular feature using trials from one condition and then testing the classifier on trials from a different condition. Above-chance cross-decoding requires that information used to make feature discriminations in the training condition is also useful to make feature discriminations in the test condition. Note that cross-decoding is improved by training on datasets with lower signal-to-noise ratio and testing on datasets with higher signal-to-noise ratio (Hurk & Beeck, 2019). Consequently, time-generalised cross-decoding was carried out by training on orientations from all imagery trials and testing on orientations from passive viewing trials. To probe whether overlap specifically related to visual processing, cross-decoding was conducted both with all 128 channels as features, and with only the posterior 56 channels as features. 128-channel time-generalised cross-decoding is shown in Figure 7, with 56-channel results shown in Appendix 1.7. In both cases, no patterns of above-chance cross-decoding were observed between real and imagined datasets, both when including all imagery trials, and when analysing only pure imagery trials. A lack of cross-decoding here suggests minimal real-imagined overlap in neural representations of orientation.

**Figure 7.**
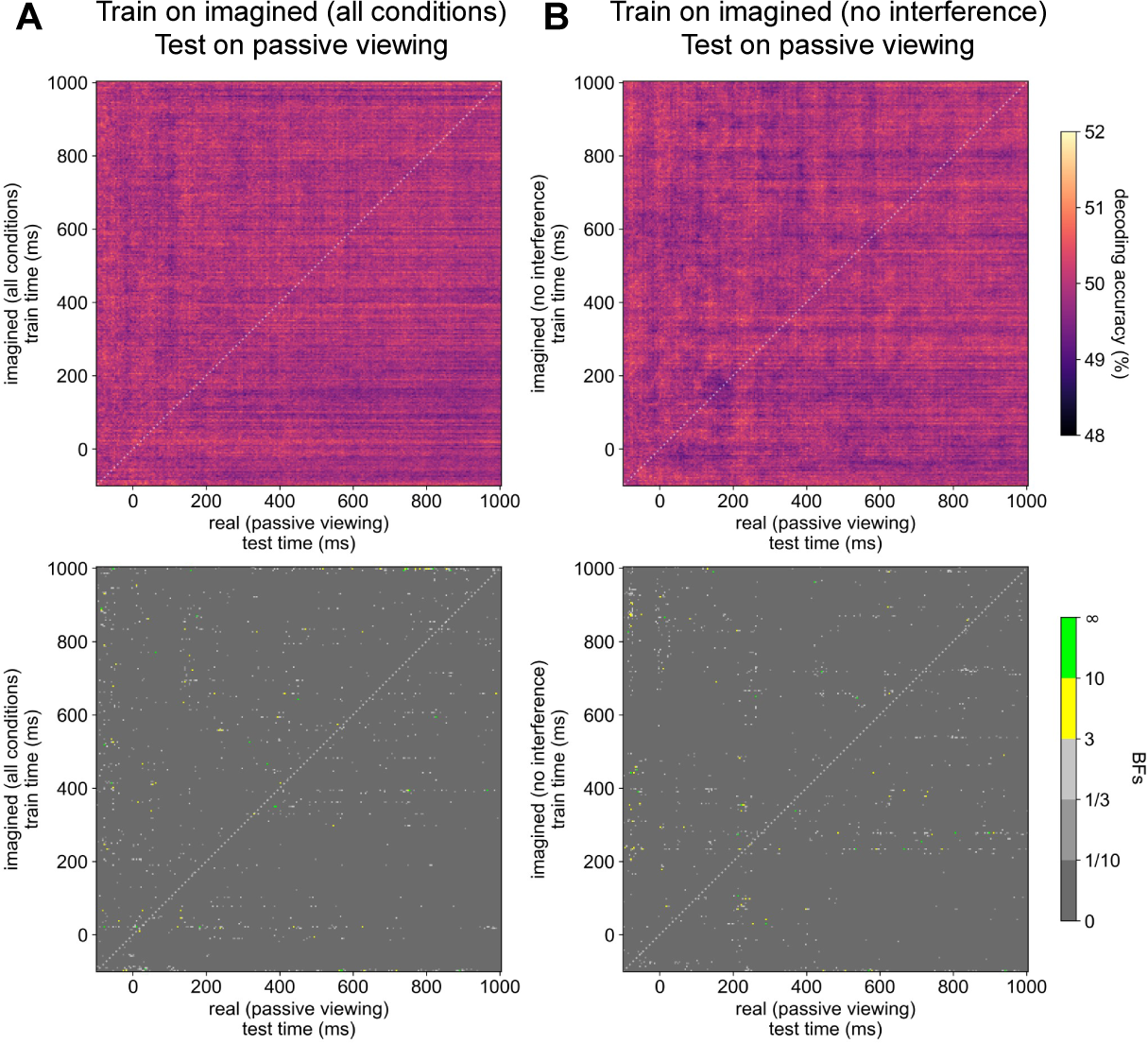
Cross-decoding: no evidence for shared representations between imagined and real orientation (128 channels) Time-generalised cross-decoding accuracy obtained using data from 128-channel EEG recordings. Classifiers were trained to discriminate imagined orientations at every timepoint during imagery trials, then tested on data from trials where participants passively viewing stimuli at the same orientations. Chance decoding accuracy is 50%. Bayes factors (BFs) are shown for the alternative hypothesis of above-chance decoding. Times are relative to image/imagery onset. A dotted white line indicates the diagonal. **A.** No pattern of time-generalised cross-decoding accuracy when training classifiers on all imagery trials and testing on passive viewing trials. **B.** No pattern of time-generalised cross-decoding accuracy when training classifiers only on imagery trials where no concurrent stimulus was presented on-screen and testing on passive viewing trials. See Appendix 1.7 for 56-channel analyses using only posterior electrodes.

Cross-decoding was also carried out between interference trials and passive viewing trials, training on the former and testing on the latter. Figure 8 shows findings for the 128-channel dataset. An expected overlap in the neural representations containing information about real orientation was observed, although a congruency-dependent effect was also observed. Neural activity patterns encoding stimulus information around ∼130–150ms during passive viewing tended to persist throughout the imagery period during interference trials, but only when real and imagined orientations were congruent. Virtually no persistence or reactivation of activity patterns was observed for the low-congruency cross-decoding condition. A similar pattern of findings was observed when analysing the posterior 56-channel only (Appendix 1.8). Differences in cross-decoding between the low- and high-congruency conditions were most pronounced at ∼148ms after stimulus onset for the 128-channel analysis. The difference in how congruency prolongs orientation representations at this specific timepoint is shown in Figure 9B & D.

**Figure 8.**
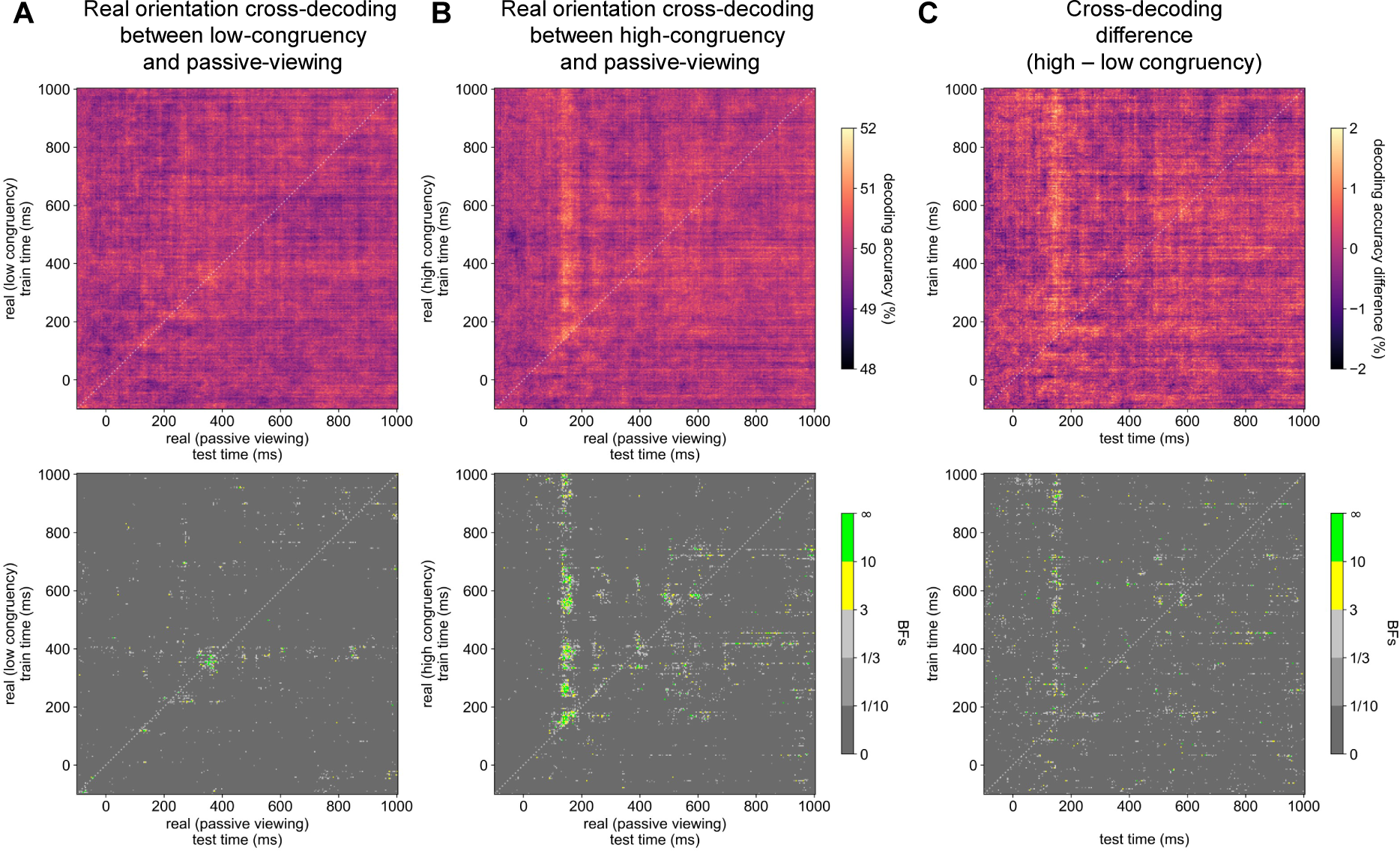
Cross-decoding: high real-imagined stimulus congruency reactivates perceptual representations (128 channels) Time-generalised cross-decoding accuracy obtained using data from 128-channel EEG recordings. Classifiers were trained to discriminate real orientations at every timepoint during interference trials, then tested on data from trials where participants passively viewing stimuli at the same orientations. Chance decoding accuracy is 50%. Bayes factors (BFs) are shown for the alternative hypothesis of above-chance decoding. Times are relative to image onset. A dotted white line indicates the diagonal. **A.** Minimal time-generalised cross-decoding when training classifiers on real orientation during poorly congruent imagery and testing on passive viewing trials. **B.** Prolonged time-generalised cross-decoding when training classifiers on real orientation during highly congruent imagery and testing on passive viewing trials. **C.** The difference in real orientation decoding accuracy between the high- and low-congruency interference conditions, indicating highly congruent imagery reactivates perceptual representations. See Appendix 1.8 for 56-channel analyses using only posterior electrodes.

**Figure 9.**
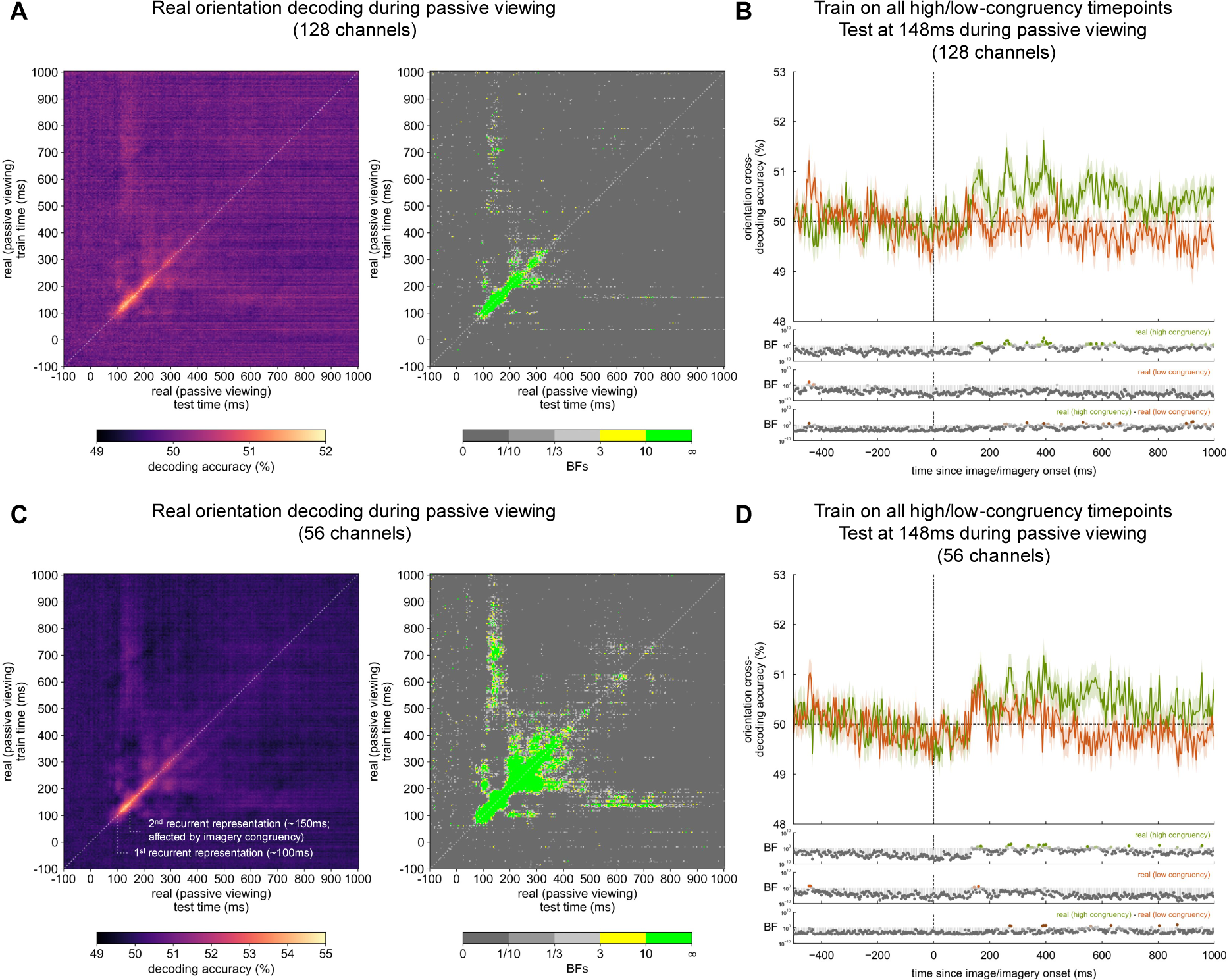
Time-generalised decoding of real orientation during passive viewing reveals the stage of perceptual processing targeted by imagery. The multi-stage representational structure of real orientation encoding. **A & C.** Time-generalised orientation decoding during passive viewing trials, showing at least two recurrent processing stages: one at ∼100ms after stimulus onset, and another at ∼150ms after stimulus onset. Chance decoding accuracy is 50%. Bayes factors (BFs) are shown for the alternative hypothesis of above-chance decoding. Times are relative to image onset. A dotted white line indicates the diagonal. **A.** Decoding using data from all 128 EEG channels. **C.** Decoding using data from only the posterior 56 EEG channels. Annotations indicate early recurrent stages, which are more visible on the 56-channel analysis. **B & D.** Real orientation cross-decoding accuracy for classifiers trained on real orientations presented during either low- or high-congruency mental imagery. Classifiers were trained on data from all timepoints during interference trials, and tested only on data from 148ms after stimulus onset during passive viewing trials as congruency-dependent differences in cross-decoding were most pronounced at this timepoint. **B.** Decoding using data from all 128 EEG channels. **D.** Decoding using data from only the posterior 56 EEG channels.

To precisely identify the activity patterns targeted by this congruency-dependent interference effect, time-generalised decoding was carried out for passively viewed orientations, on both the 128- and 56-channel datasets. Results are displayed in Figure 9A & C, showing multiple activations and reactivations of neural representations across time for externally driven perception. At least three distinct activity patterns encoding orientation were identified: an initial non-recurrent activity pattern peaking prior to 100ms after stimulus onset, then a recurrent activity pattern reactivated most at ∼100ms, followed by a second recurrent pattern reactivated at ∼150ms. These processing stages were evident in both the 128- and 56-channel analyses, but were most clearly visible in the 56-channel analysis. Imagery congruency appeared to modulate the latter recurrent representation, and not the earlier recurrent or non-recurrent representations.

## Discussion

### The contents of imagined stimulus representations

Overall, this study shows that imagination can involve information about basic visual features, and that such features can be decoded from neural activity. Neural representations of imagined content were also shown to interact with those of real perceptual content in a generally constructive manner. This study also showcased how rhythmic cues can be used to aid in time-locking instances of mental imagery, and that the deliberate construction of mental images invoked stimulus-specific neural activity patterns which could not be attributed merely to the recent encoding of a stimulus into memory. Yet, while interactions with basic visual information were observed in this study, the amount of information in the brain concerning imagined orientation was generally low, especially when compared to the information available to differentiate the orientation of real images.

The relatively low informational capacity of imagined neural representations could be due to limitations inherent to the process of mental imagery. There is a good scientific basis for imagined neural representations to have a relatively small dependence on the early visual regions which support basic visual feature representations generally. Koenig-Robert & Pearson (2021) suggest that mental imagery should have the capacity to modulate, but not drive, activity in early visual cortices. Breedlove et al. (2020) note that information loss during feedforward memory encoding means that imagined stimulus representations fed back to the visual cortex should be deficient relative to during veridical perception. Finally, mental imagery seems unaffected by low-level cortical damage, including cortical blindness (Bridge et al., 2012; Chatterjee & Southwood, 1995; de Gelder et al., 2015; Zago et al., 2010), nor mid-level damage, such as in agnosia (Behrmann et al., 1994). This implies that normal imagery processes may have little impact on low-level visual processes generally. Hence, it may not be surprising that a classifier might have little information to draw from when attempting to decode orientation, a basic visual feature, only from neural representations of imagined content.

### Interactions between real and imagined perceptual content

Although neural representations were only mildly informative in isolation, the effects of imagination were still observed via interference between real and imagined perceptual content. Real orientation seemed to be more easily decoded in the presence of highly congruent, rather than poorly congruent, mental imagery. This trend is largely consistent with a constructive-type interaction effect between real and imagined content, although a statistically significant signal amplification from congruent imagery was not observed consistently enough to make a firm conclusion. However, imagery nevertheless did seem to interact with real stimulus representations, boosting and extending activity patterns of real stimuli when they aligned with what was imagined. Future studies could systematically modulate the onset and duration of interference for a more detailed temporal analysis of interference effects.

Given that evidence for interference between real and imagined stimuli was detected here, we may expect that, like in previous studies, imagination and real perception should rely on the same neural representations of perceptual content (Albers et al., 2013; Bosch et al., 2014; Cichy et al., 2012; Dijkstra et al., 2017, 2018, 2019, 2020; Lee et al., 2012; Naselaris et al., 2015; Xie et al., 2020). However, this study found that neural representations relevant for real orientation classification during passive viewing were not relevant for classification of imagined orientation and vice versa. This suggests a lack of evidence for shared representations between mental imagery and veridical perception, at least with respect to orientation decoding. A plausible explanation is that mental images affect neural representations used during veridical perception yet lack the capacity to induce the same level of activity as real percepts. This view is supported by the finding that imagery did modulate real orientation decoding even though imagined orientation itself was not possible to decode here. Highly congruent imagery appeared to boost the persistence of real perceptual representations recruited approximately 130–150ms after stimulus onset, while poorly congruent imagery did not. This suggests that imagery is carrying stimulus-specific sensory information to neurons involved in veridical orientation perception, but not enough to induce an independent instantiation of perceptual content. The effects of imagery on basic feature perception therefore seem to be only modulatory, at least with respect to orientation, consistent with the conception of imagery suggested by Koenig-Robert & Pearson (2021).

### Mental imagery targets later stages of visual processing

The findings obtained here can be used to infer the stage of perceptual processing targeted by mental imagery. Time-generalised decoding of real orientation during passive viewing revealed distinct and recurrent neural representations of orientation, with the first recurrent neural representation beginning to reactivate at approximately 100ms after stimulus onset, while the second recurrent neural representation is activated roughly 50ms later. Perceptual processing commonly involves recurrence, with feedforward-feedback sweeps between early- and late-stage perceptual representations oscillating in the alpha frequency band (Dijkstra et al., 2020). Such an oscillation would be consistent with the pattern observed here, with feedforward processing from the first recurring activity pattern in Figure 9 to the second, and feedback processing from the second recurring activity pattern back to the first, and so on, at approximately 10Hz. Imagery congruency effects seem to specifically target the second recurrent neural representation activated during veridical perceptual processing while the earliest recurrent representation was unaffected. These findings could suggest that imagery is mainly influencing the later perceptual processing stages from which feedback signals originate during veridical perception, with no influence persisting to earlier stages.

### Considerations and limitations

Although imagined orientation was possible to decode and real-imagined congruency effects were observed, effects were generally small. It is difficult to separate whether this is because of the nature of neural processing or because of limitations in the current study. Idiosyncrasies of the experimental design used here may have limited the degree to which imagery appears to influence perceptual representations. Experimentally, this study prioritised precise imagery timing and the use of a valid unimagined control orientation at the expense of trial brevity, limiting the total number of samples of imagery available to train a classifier. Secondly, even with a rhythmic cue, precisely timed imagery is still likely to be less temporally precise than real image perception. Imagery timing might also vary between participants, with some aiming to have imagery constructed by the time the imagery cue appears on-screen, while others may only start creating imagery when the cue appears. Thirdly, imagined stimuli were always at unconventional orientations, while real bars presented on-screen during imagery were either cardinal, or intercardinal obliques. Given the immediate recognisability of cardinal and intercardinal orientations, imagined orientations were deliberately unconventional to encourage a visual mental imagery strategy during recall rather than a verbal labelling strategy. However, cardinal angles are afforded visual processing advantages relative to other angles (Appelle, 1972; Nasr & Tootell, 2012), which could add to the disparity between imagined and real orientation decoding during interference trials. Collectively, these practical considerations could reduce the likelihood of above-chance decoding of imagined orientation at any given timepoint, as well as generalisation between real and imagined representations of orientation. However, congruency effects should not be affected.

As with all mental imagery studies, it is not possible to verify whether a subjective visual experience (i.e. an actual mental image) truly occurred during the designated mental imagery period. Even though participants were instructed to conduct mental imagery during the designated period, participants could still have passed the memory checks in the study by using a working memory strategy that did not rely on mental imagery. As participants received feedback regarding their recall accuracy, they may have even been incentivised to adopt a non-imagery memory strategy if they found visualisation difficult or inaccurate. However, previous research has shown that very few people consistently report being unable to visualise simple stimuli (Sulfaro et al., 2024). Additionally, congruency-specific effects on recall accuracy were still observed, suggesting at least that participants were unlikely to be relying solely on a verbal labelling strategy for recall.

## Conclusion

Overall, this study showed that neural representations of real and imagined stimuli can affect each other in a content-specific manner consistent with a mild, constructive-type interaction. This study also showed that a rhythmic imagery paradigm can support the temporal precision of mental imagery generation such that basic imagined stimulus features can be decoded directly from EEG recordings, improving on previous EEG studies which did not report imagery decoding during the imagery period (Linde-Domingo et al., 2019; Shatek et al., 2019). Ultimately, the consequences of imagination were here best observed indirectly via mental imagery’s modulatory effect on veridical perception, with evidence collectively suggesting that mental imagery has only a limited influence over perceptual representations activated during early-stage visual perception.

