## Appendix 1 for "Neural interference between real and imagined visual stimuli"

#### Appendix 1.1

*This appendix item is included as a placeholder so that the remaining Appendix 1 figure numbers for 56-channel analyses correspond to the figure numbers for 128-channel analyses in the main text.*

#### Appendix 1.2

*Decoding of real orientation, imagined orientation, and congruency type (56 channels)*

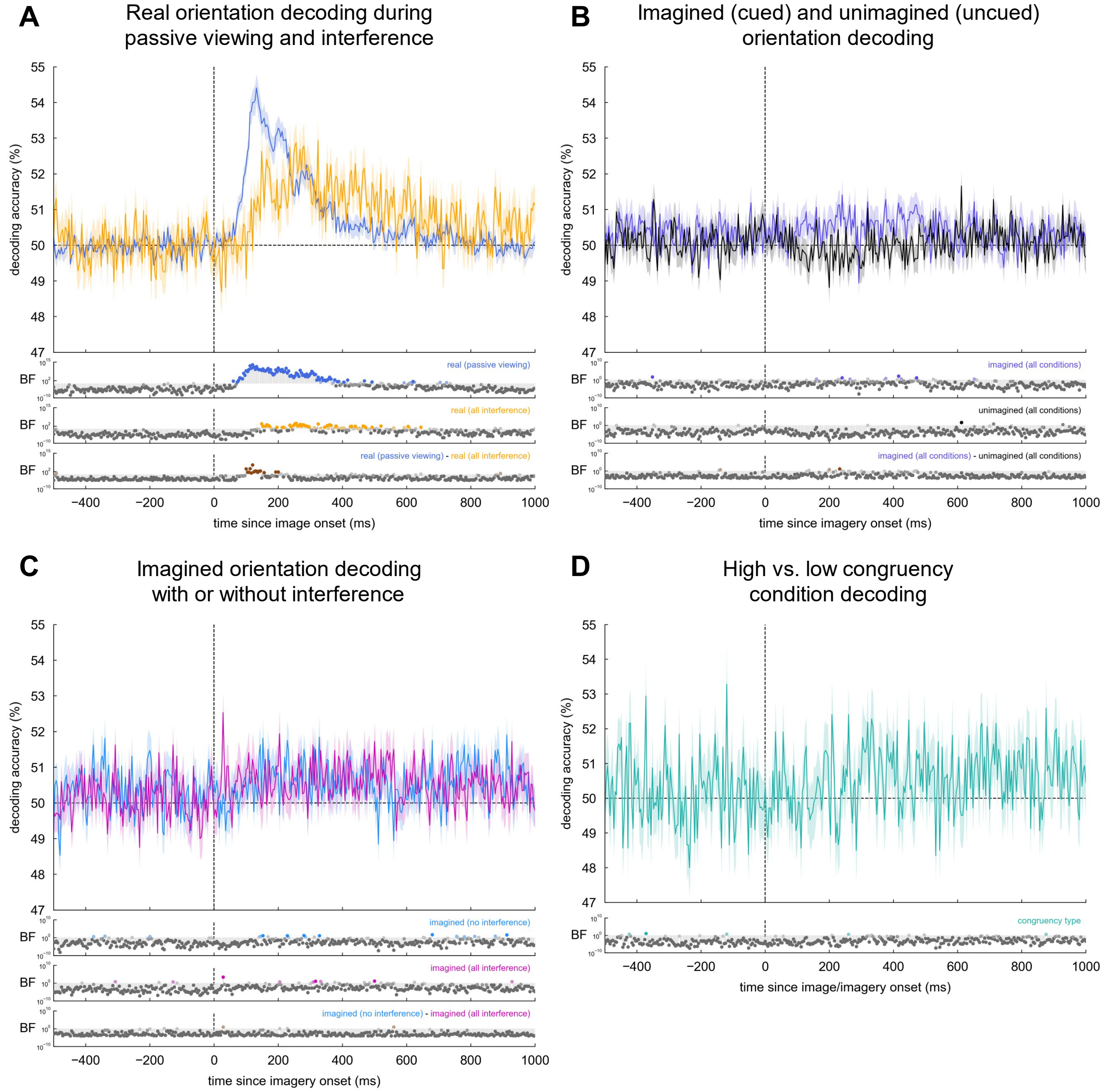


*Note.* Decoding accuracy over time from 56-channel EEG recordings. Chance decoding accuracy is 50%. Bayes factors (BF) are shown for the alternative hypothesis of above-chance decoding, or for a difference in decoding accuracy between two conditions. BF ≥ 10 (saturated colour), 3 ≤ BF < 10 (pale colour), 1/3 ≤ BF < 3 (light grey), 1/10 ≤ BF < 1/3 (grey), BF < 1/10 (dark grey). **A.** Decoding the orientation of real stimuli presented on-screen, either during passive viewing trials, or during interference trials with concurrent mental imagery. **B.** Decoding the orientation of the stimulus cued to be imagined in the imagery period (imagined), and of the stimulus which was encoded but not recalled (unimagined). **C.** Decoding imagined orientation when no competing stimulus was on-screen (no interference), or when a competing stimulus of any orientation was on-screen at the same time (all interference). **D.** Decoding whether an imagery trial was presented at the same time as a highly or poorly congruent stimulus on-screen. See Figure 2 for 128-channel analyses.

#### Appendix 1.3

*Orientation decoding during high and low congruency interference conditions (56 channels)*

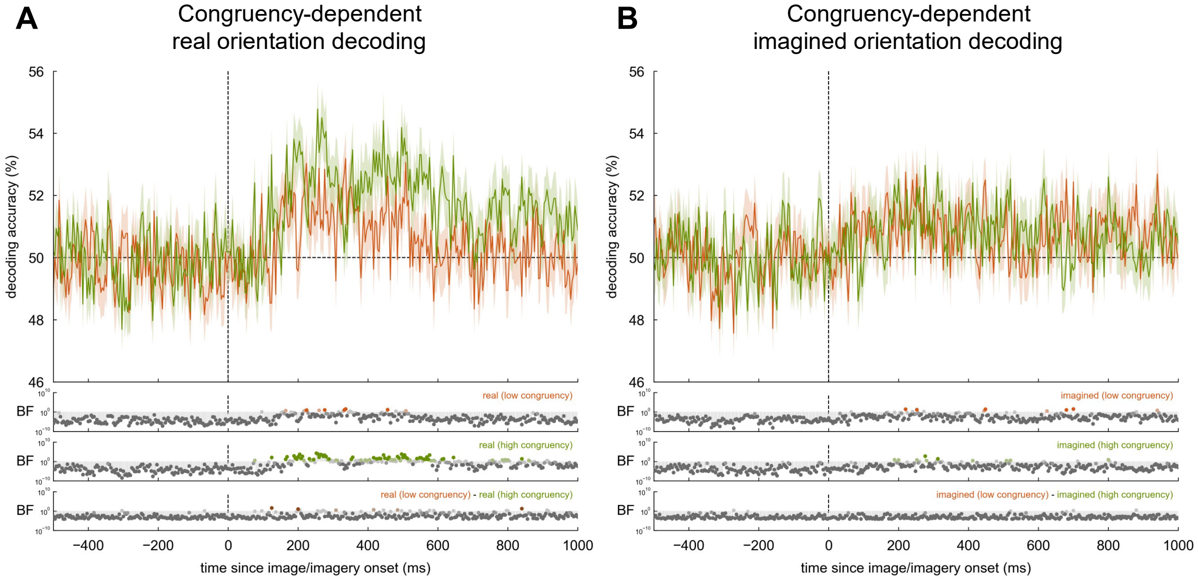


*Note.* Decoding accuracy over time from 56-channel EEG recordings. Chance decoding accuracy is 50%. Bayes factors (BF) are shown for the alternative hypothesis of above-chance decoding, or for a difference in decoding accuracy between two conditions. BF ≥ 10 (saturated colour), 3 ≤ BF < 10 (pale colour), 1/3 ≤ BF < 3 (light grey), 1/10 ≤ BF < 1/3 (grey), BF < 1/10 (dark grey). **A.** Decoding real orientation during poorly congruent mental imagery or highly congruent mental imagery. **B.** Decoding imagined orientation in the presence of a poorly congruent or highly congruent real stimulus. See Figure 3 for 128-channel analyses.

### Appendix 1.4

#### *Time-generalised decoding of imagined orientation (56 channels)*

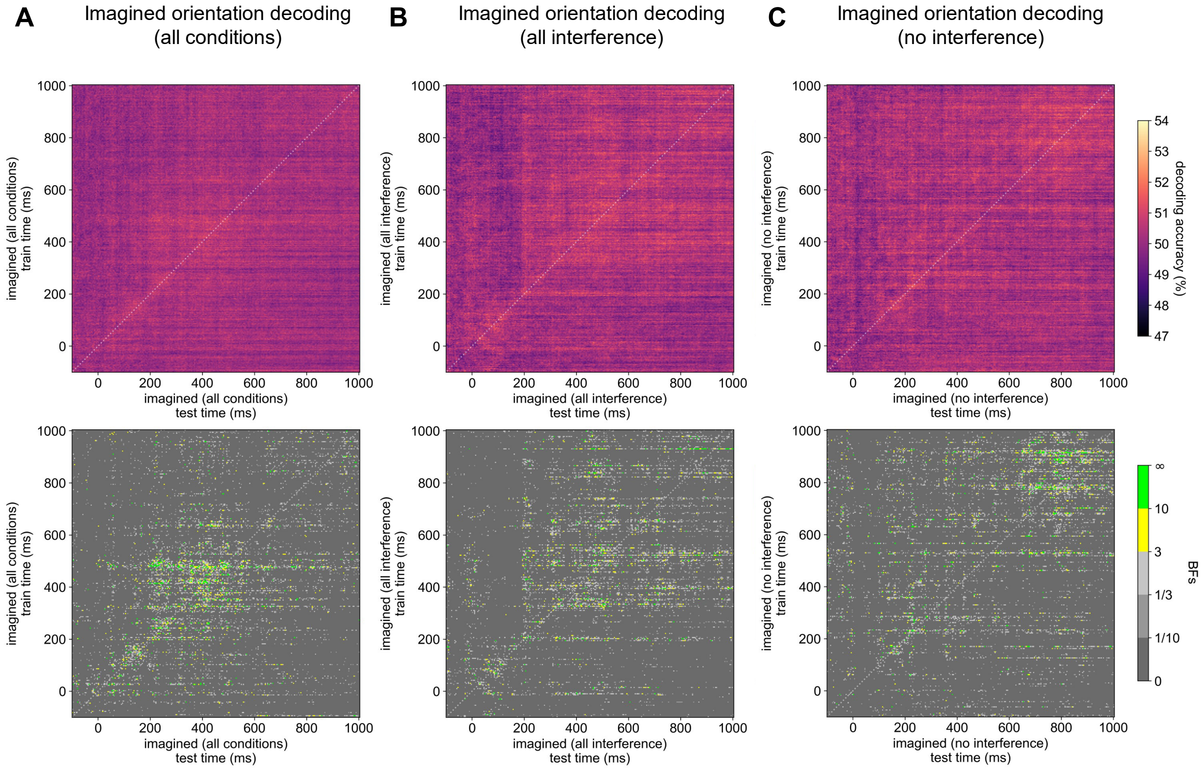


*Note.* Time-generalised decoding accuracy, training and testing a classifier at every timepoint, using data from 56-channel EEG recordings. Chance decoding accuracy is 50%. Bayes factors (BFs) are shown for the alternative hypothesis of above-chance decoding. Times are relative to imagery onset. A dotted white line indicates the diagonal. **A.** Time-generalised decoding of imagined orientation for all imagery trials. **B.** Time-generalised decoding of imagined orientation when a competing stimulus of any orientation was on-screen at the same time. **C.** Time-generalised decoding of imagined orientation when no competing stimulus was on-screen. See Figure 4 for 128-channel analyses.

### Appendix 1.5

*Time-generalised decoding of real orientation dependent on congruency condition (56 channels)*

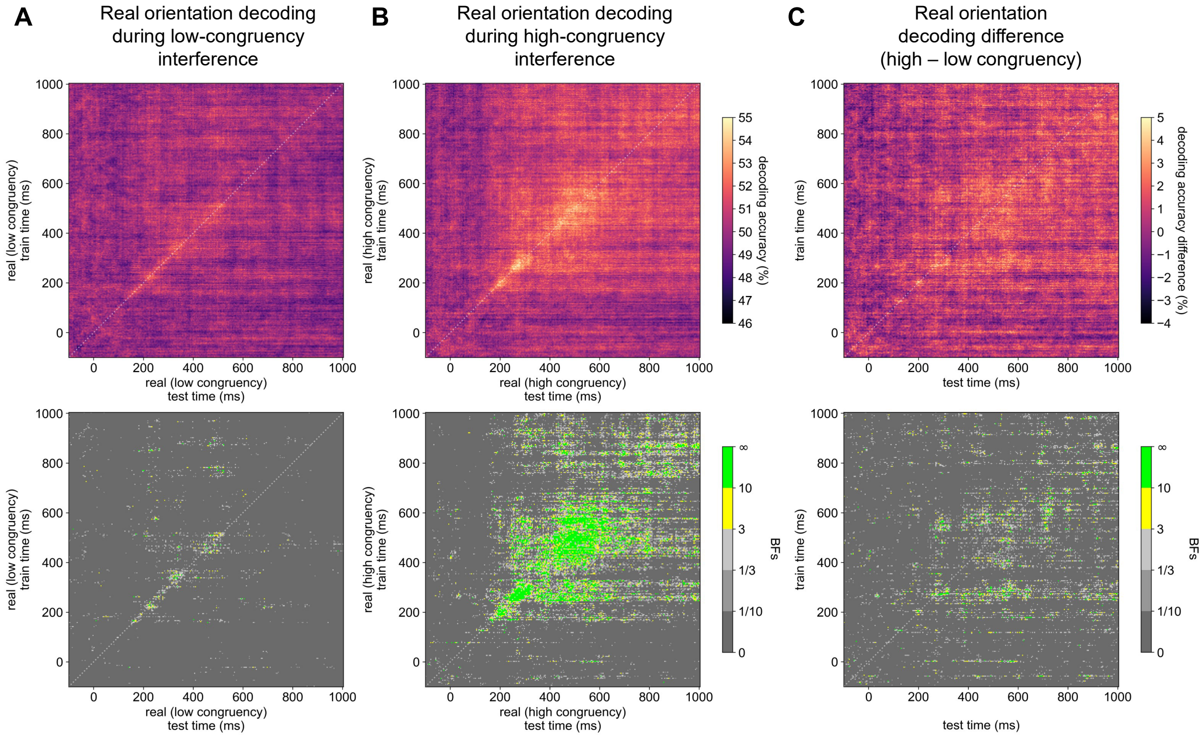


*Note.* Time-generalised decoding accuracy, training and testing a classifier at every timepoint, using data from 56-channel EEG recordings. Chance decoding accuracy is 50%. Bayes factors (BFs) are shown for the alternative hypothesis of above-chance decoding, or for a difference in decoding accuracy between two conditions. Times are relative to image onset. A dotted white line indicates the diagonal. **A.** Time-generalised decoding of real orientation during poorly congruent mental imagery. **B.** Time-generalised decoding of real orientation during highly congruent mental imagery. **C.** The difference in real orientation decoding accuracy between the high- and low-congruency interference conditions. See Figure 5 for 128-channel analyses.

### Appendix 1.6

*Time-generalised decoding of imagined orientation dependent on congruency condition (56 channels)*

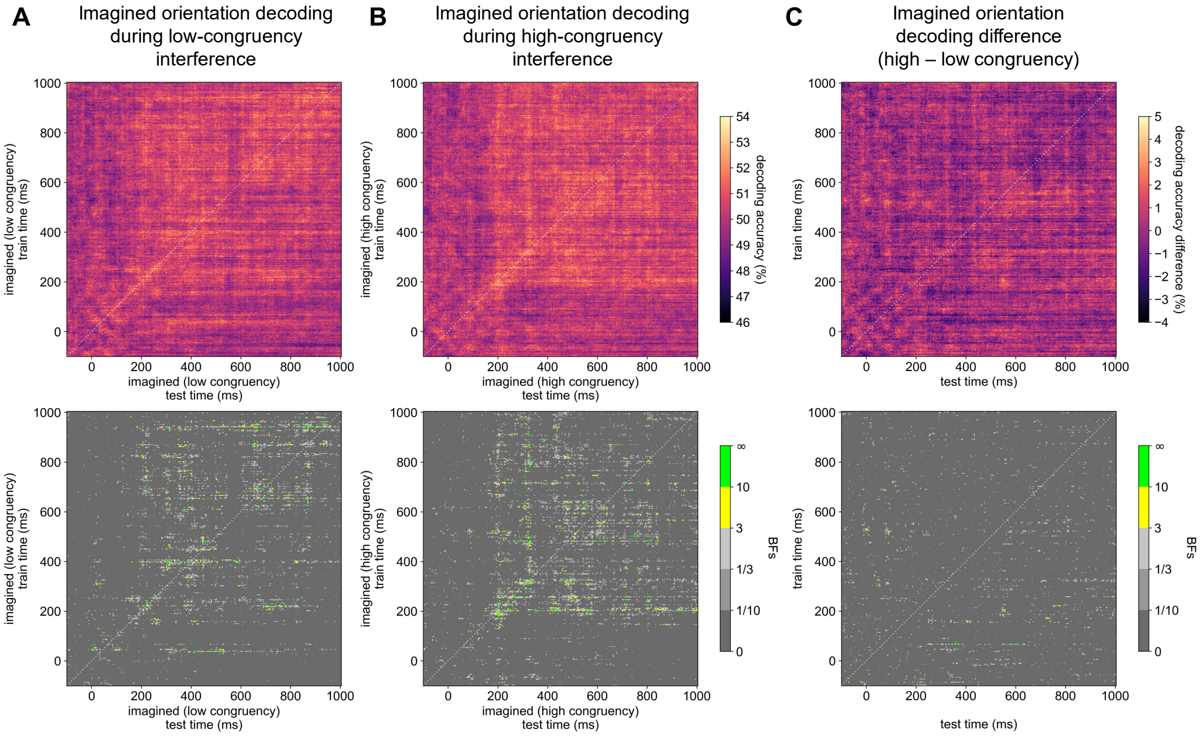


*Note.* Time-generalised decoding accuracy, training and testing a classifier at every timepoint, using data from 56-channel EEG recordings. Chance decoding accuracy is 50%. Bayes factors (BFs) are shown for the alternative hypothesis of above-chance decoding, or for a difference in decoding accuracy between two conditions. Times are relative to imagery onset. A dotted white line indicates the diagonal. **A.** Time-generalised decoding of imagined orientation in the presence of a poorly congruent real stimulus. **B.** Time-generalised decoding of imagined orientation in the presence of a highly congruent real stimulus. **C.** The difference in imagined orientation decoding accuracy between the high- and low-congruency interference conditions. See Figure 6 for 128-channel analyses.

#### Appendix 1.7

*Cross-decoding: no evidence for shared representations between imagined and real orientation (56 channels)*

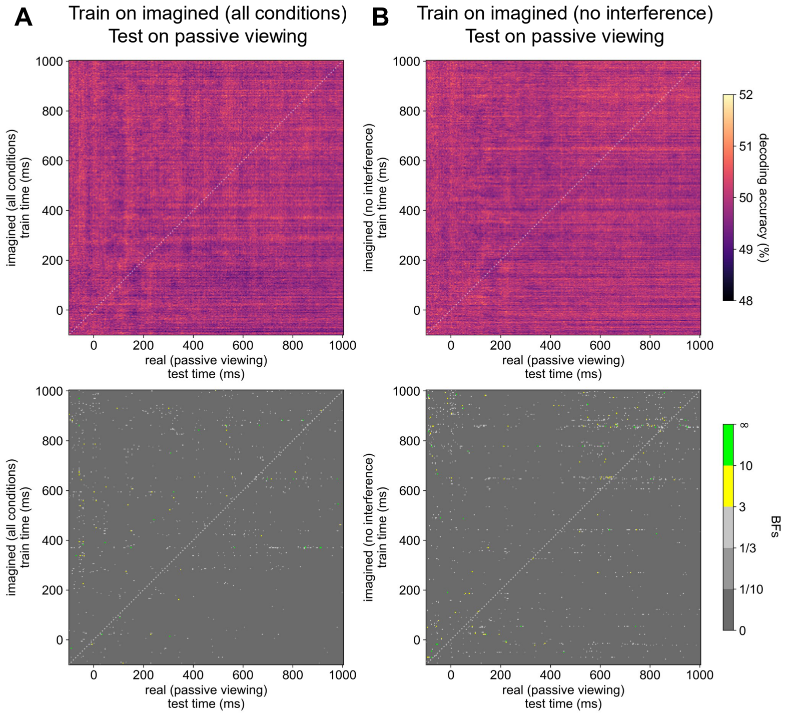


*Note.* Time-generalised cross-decoding accuracy obtained using data from 56-channel EEG recordings. Classifiers were trained to discriminate imagined orientations at every timepoint during imagery trials, then tested on data from trials where participants passively viewing stimuli at the same orientations. Chance decoding accuracy is 50%. Bayes factors (BFs) are shown for the alternative hypothesis of above-chance decoding. Times are relative to image/imagery onset. A dotted white line indicates the diagonal. **A.** No pattern of time-generalised cross-decoding accuracy when training classifiers on all imagery trials and testing on passive viewing trials. **B.** No pattern of time-generalised cross-decoding accuracy when training classifiers only on imagery trials where no concurrent stimulus was presented on-screen and testing on passive viewing trials. See Figure 7 for 128-channel analyses.

#### Appendix 1.8

*Cross-decoding: high real-imagined stimulus congruency reactivates perceptual representations (56 channels)*

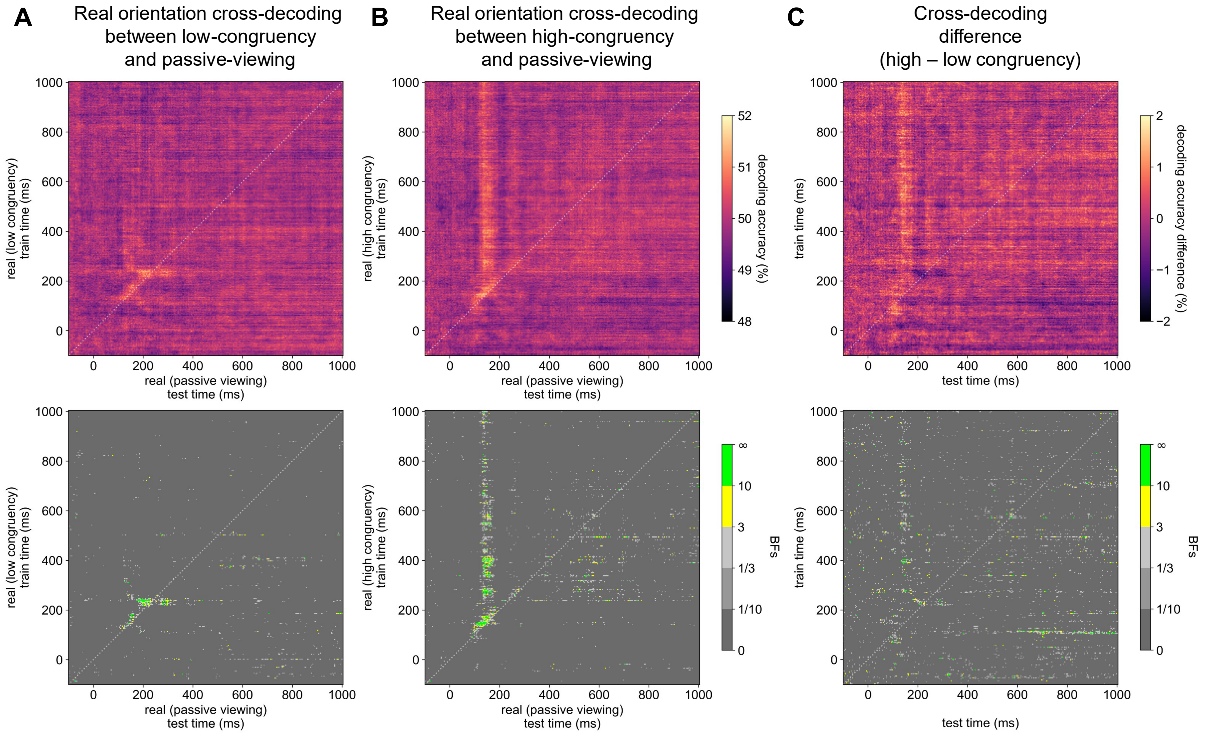


*Note.* Time-generalised cross-decoding accuracy obtained using data from 56-channel EEG recordings. Classifiers were trained to discriminate real orientations at every timepoint during interference trials, then tested on data from trials where participants passively viewing stimuli at the same orientations. Chance decoding accuracy is 50%. Bayes factors (BFs) are shown for the alternative hypothesis of above-chance decoding. Times are relative to image onset. A dotted white line indicates the diagonal. **A.** Minimal time-generalised cross-decoding when training classifiers on real orientation during poorly congruent imagery and testing on passive viewing trials. **B.** Prolonged time-generalised cross-decoding when training classifiers on real orientation during highly congruent imagery and testing on passive viewing trials. **C.** The difference in real orientation decoding accuracy between the high- and low-congruency interference conditions, indicating highly congruent imagery reactivates perceptual representations. See Figure 8 for 128-channel analyses.
